# Network topology dictates sequential drug efficacy through bistability-mediated state switching

**DOI:** 10.64898/2026.05.01.722163

**Authors:** Taha O. Osman, Karina Islas Rios, Anthony Hart, Sung-Young Shin, Lan K. Nguyen

## Abstract

Sequential drug combinations can significantly enhance therapeutic efficacy, yet the general principles governing when and why sequential administration outperforms concurrent treatment remain poorly understood. While empirical evidence demonstrates that the order and timing of drug exposure can be critical, a mechanistic framework to predict which regulatory architectures are primed for sequential benefit is currently lacking. Here, we systematically enumerated and dynamically analysed 59,040 four-node network topologies to identify the structural design principles that dictate sequential efficacy. Our analysis reveals that only a small fraction of network architectures robustly confer a sequential advantage and identifies a minimal structural requirement for this benefit: a positive feedback loop between the primary drug target and its downstream oncogenic output, coupled with antagonistic crosstalk from a secondary drug target. We demonstrate that this architecture enables bistability, allowing the first drug to reconfigure the network into a suppressed attractor state that is inaccessible through concurrent administration. The treatment schedule determines which of two coexisting stable states the system ultimately occupies, with the gap time between doses defining a critical therapeutic window. Only when the first drug is given sufficient time to displace the system past a threshold does the sequential regimen achieve superior suppression. Our findings establish bistability-enabling network motifs as predictive determinants of sequential drug efficacy and provide a topology-based framework for the rational design of time-dependent combination therapies.

## Introduction

Although major advances in cancer treatment have improved survival and patient outcomes [1], only a small proportion of patients with advanced-stage disease achieve long-term remission or cure, with cure rates falling to single digits in most cancers [2]. Tumour relapse and the development of therapy resistance are major contributors to these poor outcomes [3]. Resistance is often driven by the complexity of tumour biology, including the dysregulation of multiple signalling pathways and extensive tumour heterogeneity, which allows different subclones to survive treatment through compensatory mechanisms [4,5]. These factors limit the effectiveness of single-agent therapies and make durable responses difficult to achieve.

Combination therapies have thus become a central strategy in modern oncology [6]. By targeting complementary signalling routes, such strategies aim to improve efficacy, suppress resistant subclones, and increase the likelihood of long-term remission. In practice, most combinations are administered concurrently, an approach that has demonstrated clinical benefit across multiple cancer types [7–9]. However, concurrent administration implicitly assumes that the therapeutic effect of a drug combination is independent of the timing of each drug’s delivery - an assumption that may not hold in the context of highly dynamic and nonlinear signalling networks.

Increasing evidence indicates that the order and timing of drug administration can critically influence therapeutic outcome [10,11]. Sequential drug exposure can produce markedly different, and in some cases superior, responses compared to simultaneous treatment. For example, Lee et al. demonstrated that time-staggered inhibition of EGFR signalling prior to DNA-damaging chemotherapy dramatically enhanced apoptosis in triple-negative breast cancer cells, an effect that was abolished under concurrent administration [12]. Similarly, Salvador-Barbero et al. showed that sequential application of CDK4/6 inhibitors after taxane-based chemotherapy cooperated to suppress tumour cell proliferation in pancreatic cancer models, whereas concurrent treatment did not, owing to the requirement for cells to progress through the cell cycle before CDK4/6 inhibition could prevent recovery from chromosomal damage [13]. Beyond targeted therapy-chemotherapy combinations, sequential scheduling has also been shown to enhance efficacy in targeted therapy pairs, as demonstrated by the sequential administration of sorafenib and sunitinib in renal cell carcinoma [14]. Despite these compelling examples, the findings have remained context-specific, with limited understanding of the general principles that govern when and why sequential treatment outperforms concurrent administration.

A common thread across these studies is that sequential efficacy arises from dynamic rewiring of signalling networks: prior drug exposure alters the cellular state and reshapes downstream responses to subsequent perturbations. This observation suggests that the capacity for sequential advantage may be encoded not in the drugs themselves but in the topology of the regulatory networks they target. If so, the question becomes: what structural features of a signalling network make it amenable to time-dependent therapeutic intervention?

Computational systems biology has demonstrated that network topology plays a critical role in determining cellular responses to perturbations. Network motifs - recurring patterns of interactions such as feedback and feedforward loops - have been identified as fundamental building blocks that encode distinct dynamical behaviours [15]. Systematic enumeration of three-node circuits has revealed that only a small number of topologies are capable of robust biochemical adaptation [16], establishing the principle that complex cellular behaviours can be predicted from network architecture. Similarly, motif-based analyses of enzymatic networks have shown that network structure can determine whether drug combinations exhibit synergy or antagonism [17], directly linking topology to therapeutic outcome. More broadly, positive feedback loops have been identified as essential structural elements for generating bistability and switch-like behaviour in biological systems [18–22], and bistable networks have been implicated in cell-fate decisions, signalling memory, and the emergence of drug-resistant states [22–24]. Together, these studies demonstrate that complex cellular behaviours can be understood and predicted from underlying network architecture.

Motivated by these observations, we hypothesised that the benefit of sequential therapy may also be governed by underlying network topology. Specifically, we propose that there exist generalisable network motifs and architectural features that confer sensitivity to drug order and timing, thereby enabling sequential treatment to outperform concurrent administration. To test this, here we systematically analysed regulatory network topologies using a large-scale *in silico* modelling and massive simulation framework, quantifying their propensity to exhibit a sequential advantage across diverse parameter regimes. Our analysis reveals that sequential benefit is not a generic property of combination therapy, but an emergent property of specific network architectures - in particular, positive feedback motifs and the bistable dynamics they enable.

These findings provide a unifying, topology-based framework for understanding temporal effects in combination therapy and offer a mechanistic foundation for the rational design of time-dependent treatment strategies. By linking network architecture to sequential treatment sensitivity, this work provides guiding principles for identifying the cancer signalling contexts and target combinations most likely to benefit from temporally structured dosing.

## Results

### A computational framework for evaluating sequential combination therapy

To identify network architectures that confer a sequential therapeutic advantage, we established a computational framework to systematically enumerate and analyse four-node regulatory networks. We reasoned that a minimal architecture capable of representing two drug-targeted pathways and their potential crosstalk is essential for capturing the functional properties of combination treatments without introducing complexity that would hinder analysis. Accordingly, each network comprises four nodes organised into two parallel pathways: X₁ and X₃ form the first pathway, and X₂ and X₄ form the second, where X₁ and X₂ represent upstream drug targets and X₃ and X₄ represent downstream oncogenic outputs (**Figure 1A**). The fixed activating edges X₁→X₃ and X₂→X₄ define these pathway axes, while all remaining inter- and intra-pathway connections are allowed to vary.

**Figure 1.**
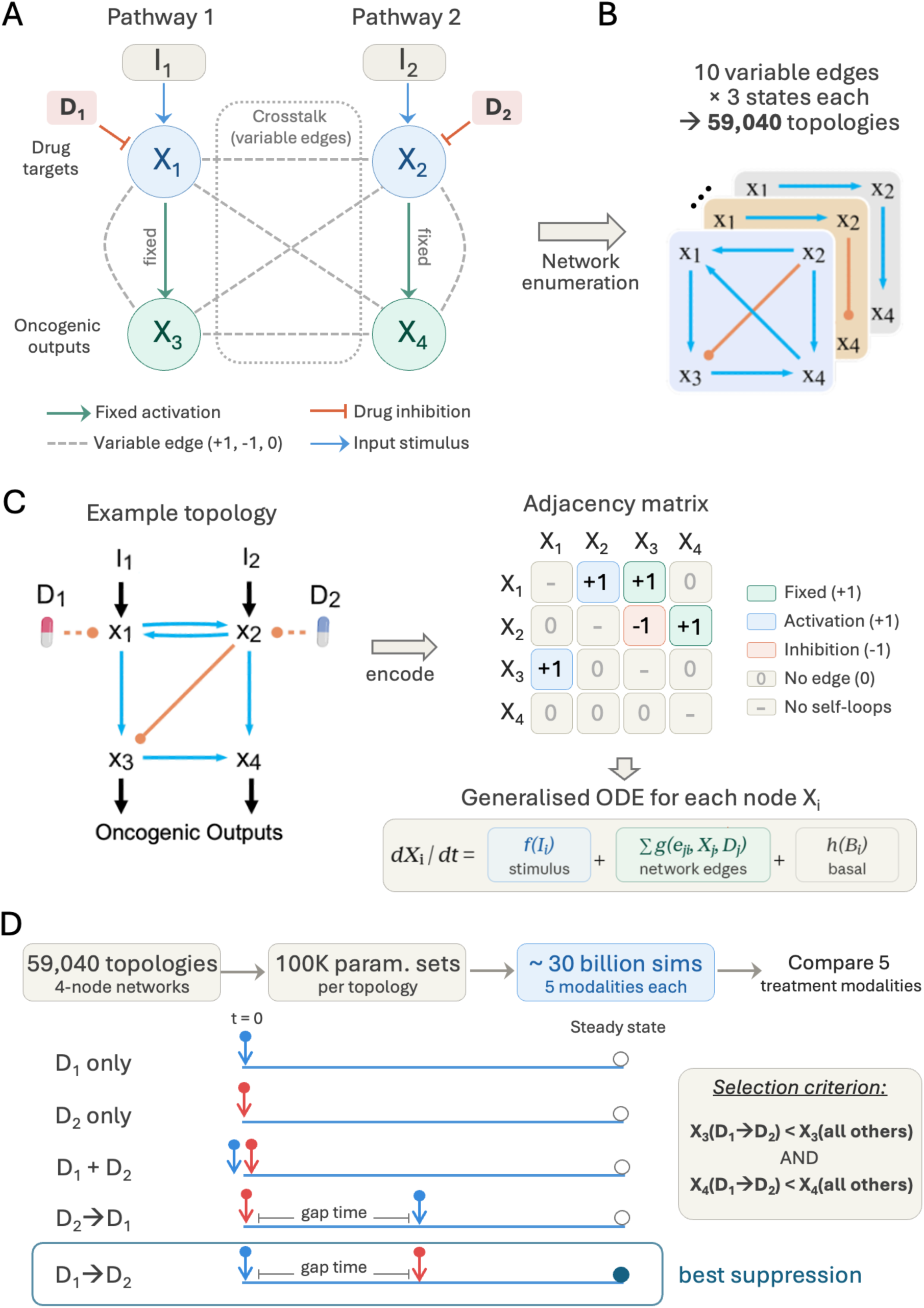
A computational framework for evaluating sequential combination therapy. **(A)** Schematic of the four-node regulatory network architecture. Nodes X₁ and X₂ represent upstream drug targets and X₃ and X₄ represent downstream oncogenic outputs, organised into two parallel pathways (X₁→X₃ and X₂→X₄). The within-pathway activations (green arrows) are fixed, while all remaining inter- and intra-pathway edges (dashed grey lines) are variable, each taking a value of +1 (activating), −1 (inhibiting), or 0 (absent). Drug inhibitions D₁ and D₂ act on X₁ and X₂, respectively, and external input stimuli I₁ and I₂ drive network activation. **(B)** Network enumeration. With 10 variable edges and 3 possible states each, the framework generates 59,040 candidate topologies after filtering for biological relevance (exclusion of self-loops, fixation of within-pathway edges, and requirement for at least one crosstalk edge). **(C)** An example topology and its mathematical encoding. The network is represented as a 4×4 adjacency matrix, from which a generalised system of ODEs based on Michaelis–Menten kinetics is constructed. Each node’s dynamics are governed by input stimulation, network-based regulation (incorporating drug inhibition via allosteric suppression), and basal activation/inhibition terms. Quality-control filters ensure that simulations converge to stable steady states within biologically meaningful bounds. **(D)** Treatment modality comparison. For each topology–parameter set combination, five treatment regimens are simulated: D₁ monotherapy, D₂ monotherapy, concurrent combination (D₁ + D₂), and two sequential schedules (D₁→D₂ and D₂→D₁), where the gap time defines the interval between drug administrations. A topology–parameter set combination is classified as conferring a sequential advantage (termed a sequential set) if the D₁→D₂ regimen achieves lower steady-state levels of both X₃ and X₄ compared to all other treatment modalities.

Each variable edge can take one of three states - activating (+1), inhibiting (−1), or absent (0) - yielding 3^16^ possible configurations in principle. To focus on biologically relevant architectures, we excluded self-regulatory interactions, fixed the within-pathway activations (X₁→X₃ and X₂→X₄), and required at least one crosstalk edge between pathways, producing a refined ensemble of 59,040 network topologies for systematic investigation (**Figure 1B**; **Methods - Network Enumeration**).

The dynamic behaviour of each topology was modelled using a system of ordinary differential equations (ODEs) based on Michaelis–Menten kinetics, in which each node’s activity is governed by input stimulation, network-based regulation, and basal turnover (**Figure 1C**; **Methods - Mathematical Model of Network Topologies**). Drug inhibition is modelled as allosteric suppression of the catalytic output of the targeted node, with the timing of administration controlled by a Heaviside step function.

To comprehensively explore the range of dynamic behaviours accessible to each topology, we generated 100,000 kinetic parameter sets using Latin Hypercube Sampling and simulated each topology across all parameter sets. Simulations were run to steady state, and only those satisfying stability and boundedness criteria were retained for downstream analysis (**Methods - Simulation Framework).**

We evaluated five treatment regimens for each topology–parameter set combination: monotherapy with drug D₁ (targeting X₁), monotherapy with drug D₂ (targeting X₂), concurrent combination therapy (D₁ + D₂), and two sequential schedules: D₁ followed by D₂ (D₁→D₂) and D₂ followed by D₁ (D₂→D₁). In each sequential regimen, the first drug is administered and the system is allowed to evolve for a defined interval - the gap time - before the second drug is introduced. Across all topologies, parameter sets, and treatment modalities, this framework comprised approximately 30 billion individual simulations (**Figure 1D**).

A topology–parameter set combination was classified as conferring a sequential advantage if the D₁→D₂ regimen achieved lower steady-state levels of both oncogenic outputs X₃ and X₄ compared to all other treatment strategies, including concurrent administration and the reverse sequential order (**Figure 1D**). The inclusion of D₂→D₁ as a comparator ensures that any identified advantage reflects a genuine order-dependent effect rather than an artefact of a particular sequence. The propensity of each topology to confer a sequential advantage was then quantified by the fraction of parameter sets satisfying this criterion.

### Identification of minimal robust sequential topologies

Of the 59,040 topologies screened, 9,716 were capable of producing a sequential advantage for at least one parameter set. However, we sought to identify topologies for which this behaviour is not merely possible but robust, that is, consistently observed across a substantial fraction of the sampled parameter space rather than arising only under rare kinetic conditions. To assess this, we quantified the number of parameter sets conferring a sequential advantage (termed sequential sets, or S-sets) for each topology across the full ensemble (**Figure 2A**). This analysis revealed that the vast majority of topologies produce sequential behaviour only sporadically, while a small subset exhibits markedly elevated S-set counts, indicating that their capacity for sequential advantage is an intrinsic property of the network architecture rather than a parameter-specific coincidence.

**Figure 2.**
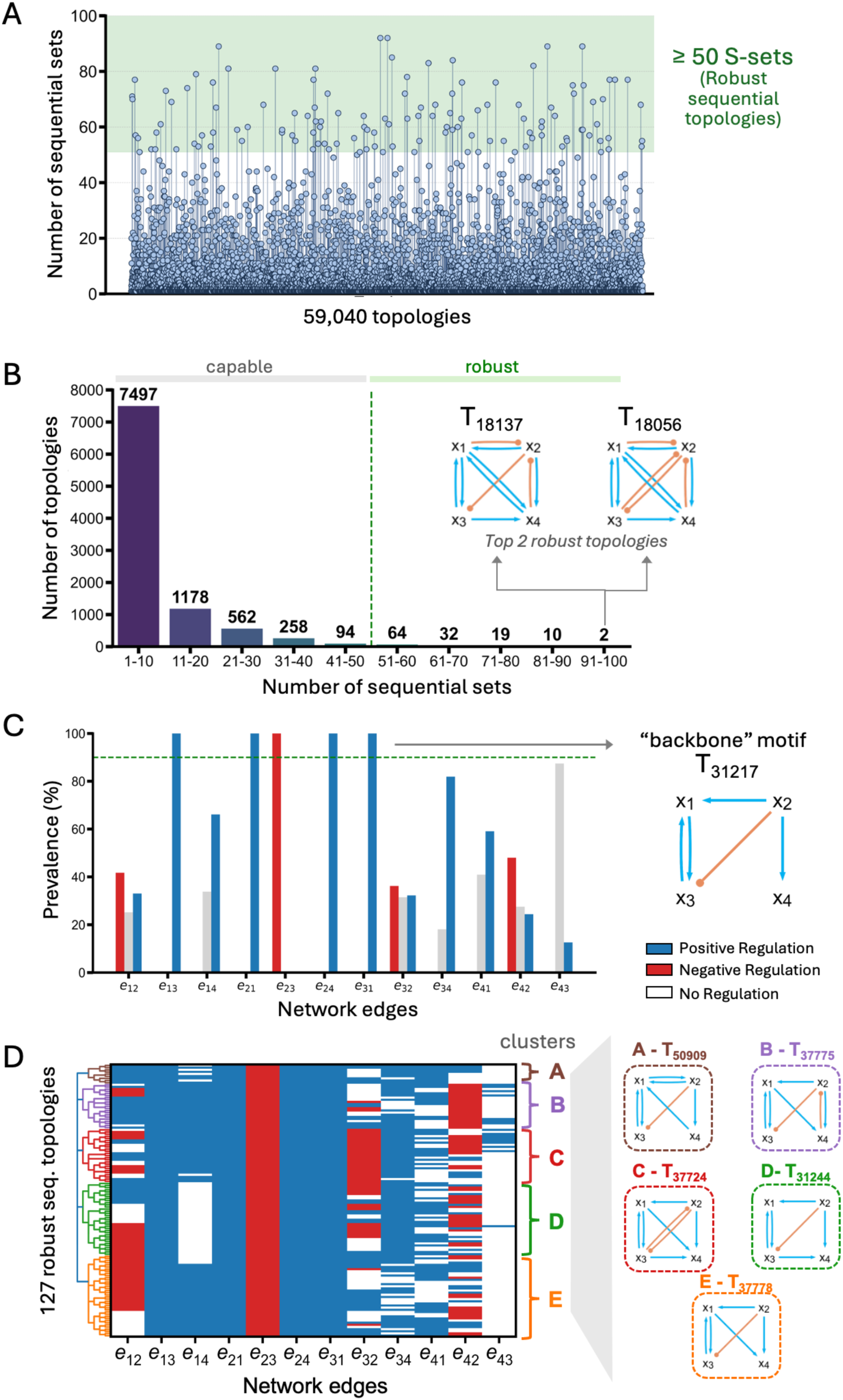
Identification of minimal robust sequential topologies. **(A)** Sequential set (S-set) frequency across the topology ensemble. Each bar represents a topology (out of 59,040), with the y-axis indicating the number of parameter sets (out of 100,000) for which the D₁→D₂ regimen achieves the best suppression. The majority of topologies produce sequential behaviour only sporadically, while a distinct subset exhibits elevated S-set counts. Topologies with more than 50 sequential sets (green shading) are classified as robustly sequential. **(B)** Distribution of topologies by sequential set frequency, binned in intervals of 10. Of 9,716 topologies capable of producing any sequential advantage, only 127 exceed the robustness threshold of 50 sequential sets. The two most robust topologies, T_18137_ and T_18056_ (92 sequential sets each), are highlighted. **(C)** Edge prevalence across the robust set, with a dashed line indicating the 90% threshold used to define the backbone topology T_31217_. The backbone comprises mutual activation between X₁ and X₃ (e₁₃, e₃₁), inhibition from X₂ to X₃ (e₂₃), and activation from X₂ to X₄ (e₂₄). Signal propagation in the e₄₃ direction is rare. **(D)** Hierarchical agglomerative clustering with a heatmap showing the state of each variable edge (blue = activating, red = inhibiting, white = absent) across all robust topologies, identifying five distinct structural families (clusters A–E) with their representative core topologies.

Based on this separation, we defined a topology as robustly sequential if it produced more than 50 sequential sets out of the total sampled parameter sets. This threshold was chosen based on a bin-to-bin retention analysis, which quantifies the proportion of topologies in each frequency bin that persist into the next higher bin. This analysis revealed a sharp inflection at approximately 50 sequential sets, above which retention increases markedly (**Supplementary Figure S1A**), indicating that topologies exceeding this threshold represent a structurally distinct, stable subset rather than the tail of a continuous distribution. By this criterion, 127 topologies qualified (**Figure 2B**), representing a highly restricted subset of architectures that robustly support sequential treatment efficacy. The two most robustly sequential topologies, T_18137_ and T_18056_, each produced 92 sequential sets.

To identify the structural features shared by these robust topologies, we analysed the prevalence of each edge across the 127 architectures (**Figure 2C**). We first asked whether a conserved backbone exists: a minimal set of interactions present in the vast majority of robust topologies. This analysis revealed that four edges occur in over 90% of the robust set: the positive feedback loop between X₁ and X₃ (edges e₁₃ and e₃₁), the inhibitory input from X₂ to X₃ (e₂₃), and the activating input from X₂ to X₄ (e₂₄). Together, these define the backbone topology T_31217_, which comprises a mutual activation between the first drug target and its oncogenic output, coupled with antagonistic crosstalk from the second drug target onto the first pathway. Notably, signal propagation in the reverse direction (e₄₃, from X₄ to X₃) is rare across the robust sets, suggesting that coordinated suppression of the two outputs is best achieved through unidirectional coupling.

Importantly, T_31217_ alone does not produce a robust sequential advantage: while it establishes the within-pathway feedback and cross-pathway antagonism required for differential drug responses, it lacks a mechanism to propagate the effect of the first drug’s perturbation to the second oncogenic output, X₄. This raised the question of what additional structural features are needed beyond the backbone.

To address this, we applied hierarchical agglomerative clustering to the 127 robust topologies, identifying five distinct structural families (clusters A–E; **Figure 2D**). Examining the edge prevalence within each cluster revealed a shared requirement across all five: the presence of at least one positive edge linking the first pathway (X₁ or X₃) to the second pathway’s oncogenic output (X₂ or X₄). In other words, signal must propagate from the first to the second pathway in order for the sequential advantage to emerge on both outputs X₃ and X_4_. This inter-pathway coupling is what enables the first drug’s action to influence the steady-state level of X₄, which the backbone alone cannot achieve.

To confirm that these structural findings are not sensitive to the specific robustness threshold, we repeated the clustering and interaction prevalence analyses at thresholds of 40 and 60 sequential sets. In both cases, the backbone architecture and cluster structures remained largely consistent (**Supplementary Figure S1B**), indicating that the identified design principles are stable properties of the sequential topology landscape rather than artefacts of a particular cutoff.

We further validated the inter-pathway coupling requirement by systematically enumerating all variants of the backbone topology T_31217_ in which positive, negative, or null connections were introduced from X₁ and X₃ to X₂ and X₄ (**Supplementary Figure S2**). Consistent with the clustering analysis, only variants with a positive inter-pathway connection produced a sequential advantage; negative or absent connections failed to propagate the first drug’s effect, eliminating the benefit. Performing the equivalent analysis for the reverse drug order (D₂→D₁) yielded structurally symmetric results - the same architectural requirements apply, but mirrored across the two pathways (**Supplementary Figure S3**) - confirming that the identified features are general properties of sequential scheduling rather than artefacts of a particular drug order.

The two largest clusters, D and E (**Figure 2D**), are defined by the core topologies T_31244_ and T_37778_, respectively. These topologies realise the inter-pathway coupling through a direct connection to X₄ - via X₃ in T_31244_ (edge e₃₄) or via X₁ in T_37778_ (edge e₁₄) - rather than through an indirect route. The prevalence of these direct connections among the most robust topologies suggests that shorter signal paths more effectively transmit the state change required for sequential benefit, a pattern we explore further through the structural analysis in the next section.

### The structural landscape of robust sequential networks

To understand how robust sequential architectures relate to one another and to the broader topology landscape, we constructed a topology atlas that maps the structural relationships among the top 500 most robust networks (**Figure 3A**). In this representation, each topology is depicted as a node and arranged into layers according to its complexity, defined by the total number of regulatory interactions. Edges connect topologies that differ by a single interaction addition or removal, thereby capturing the minimal structural transformations between neighbouring architectures (**Methods - Topology Atlas**).

**Figure 3.**
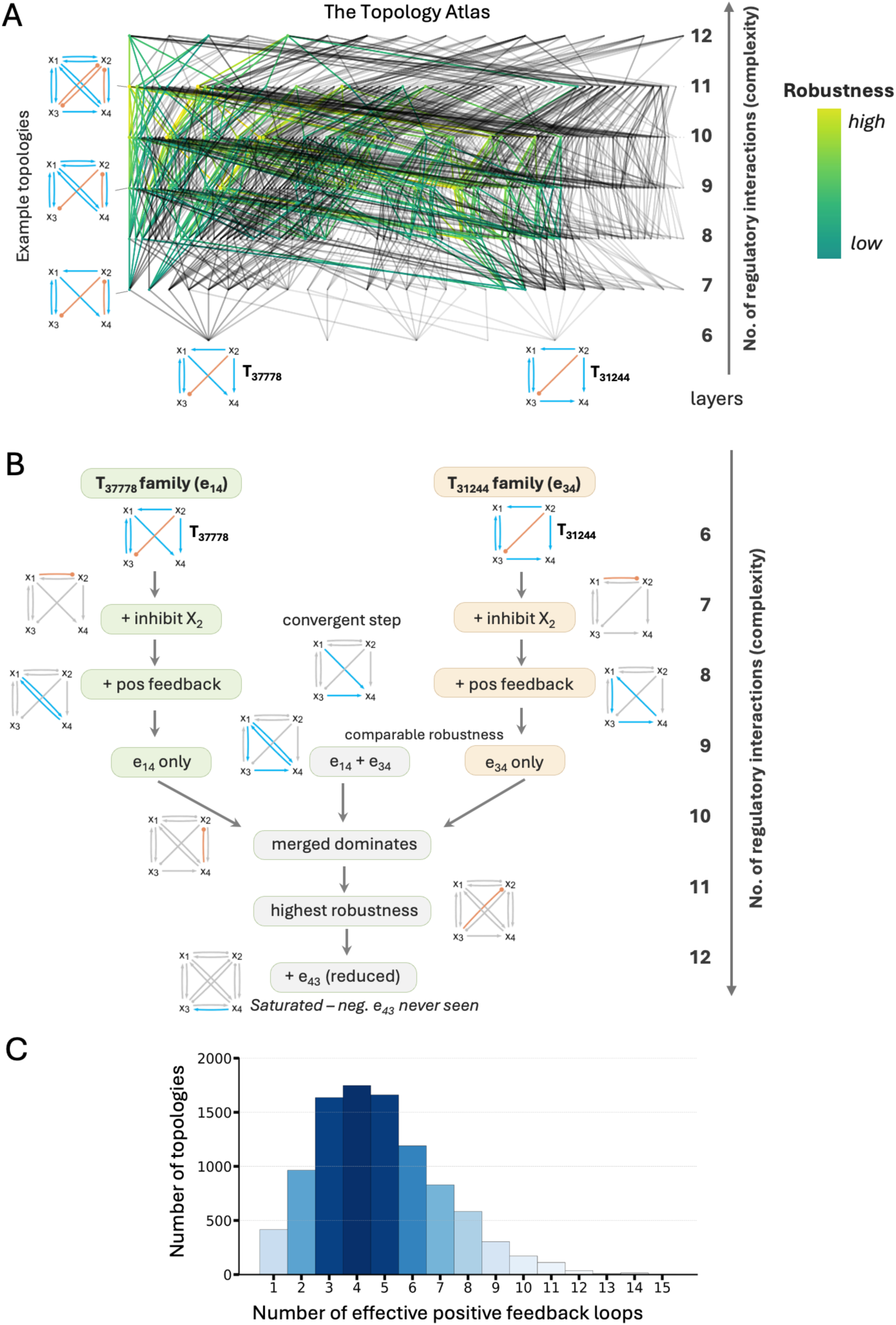
Structural landscape and design hierarchy of robust sequential networks. **(A)** Topology atlas of the 500 most robust networks, organised into layers by complexity (number of regulatory interactions). Each node represents a topology, with size and opacity scaled by robustness. Edges connect topologies differing by a single interaction addition or removal. Robust sequential topologies (coloured) descend from two dominant families defined by backbones T_37778_ (green; characterised by edge e₁₄) and T_31244_ (yellow; characterised by edge e₃₄). **(B)** Structural progression of the two families across increasing complexity (6–12 edges), showing representative topologies at each layer. At the 7-edge layer, both families independently acquire inhibition of x₂ (convergent step). Positive feedback loops accumulate progressively. By the 10-edge layer, the merged architecture (containing both e₁₄ and e₃₄) becomes dominant. At 12 edges, the addition of e₄₃ reduces robustness; negative e₄₃ is never observed. Gold, silver, and bronze labels denote the most, second most, and third most robust topologies overall. Asterisks indicate relative robustness ranking within each layer. The complete topology set is provided in **Supplementary Figure S4**. **(C)** Distribution of effective positive feedback loops (including direct positive feedback and double-negative feedback) across all 9,716 sequential topologies. Every sequential topology contains at least one effective positive feedback loop, establishing this as a universal structural requirement.

The atlas reveals that robust sequential topologies do not arise independently across the landscape but instead descend from a shared structural lineage rooted in the two core topologies identified previously: T_37778_ (characterised by edge e₁₄) and T_31244_ (characterised by edge e₃₄). This indicates that the sequential advantage is an emergent property constrained by the architectural features encoded within these backbones, with robustness progressively enhanced through the addition of further regulatory interactions.

To characterise this progressive enhancement, we traced the structural evolution of both families across increasing levels of complexity, highlighting representative topologies at each layer (**Figure 3B**; an expanded set is provided in **Supplementary Figure S4A**). At the 7-edge layer, both families independently acquire a negative regulatory interaction targeting the second drug node, X₂, revealing a shared structural step. Across all subsequent layers, the most robust topologies consistently retain this inhibition of X₂, highlighting its role as a conserved design feature. As complexity increases further, positive feedback loops are progressively incorporated into the core architectures, indicating that positive feedback reinforcement is a selectively favoured design principle for stabilising sequential behaviour.

At the 8-edge layer, the two families begin to converge; however, the most robust topologies remain those that preserve direct signal propagation from X₃ to X₄ (edge e₃₄). By the 9-edge layer, two competing architectural variants coexist with comparable robustness: one defined by the combined presence of e₁₄ and e₃₄, and another relying solely on e₃₄. This balance shifts at the 10-edge layer, where the merged architecture becomes dominant, reflecting a structural advantage conferred by enhanced coupling from X₁ to X₄. By the 11-edge layer, these features are consolidated, with the merged architecture dominating the landscape and exhibiting the highest robustness.

From this progression, a hierarchy of structural priorities emerges for enhancing robustness beyond the core sequential backbone. First, direct signal propagation from the primary oncogenic output X₃ to the secondary output X₄ is consistently preserved as a core requirement. Second, suppression of X₂ - mediated either by X₁ or by X₄ - is strongly favoured. Third, the accumulation of positive feedback loops serves as a reinforcing mechanism that stabilises the desired dynamics. Finally, indirect signal propagation from X₁ to X₄ provides an additional, albeit secondary, structural refinement.

At maximal complexity (12 edges), the network effectively saturates these design principles. The only remaining edge to be added, e₄₃, introduces reverse signal propagation from X₄ to X₃. While a positive e₄₃ can be accommodated, it consistently reduces robustness (**Figure 3A-B**). A negative e₄₃ interaction is never observed among robust topologies, consistent with the requirement for coordinated suppression of both oncogenic outputs: a negative coupling would impose opposing dynamics between X₃ and X₄ and therefore be detrimental to sequential benefit.

The recurrent appearance of positive feedback loops throughout this structural hierarchy prompted us to examine whether this feature is universal across all sequential topologies, not just the robust subset. Using effective feedback loop analysis **(Methods - Effective Positive Feedback Loop Analysis)**, we quantified the number of positive feedback structures, including both direct positive feedback and double-negative feedback loops, present in each of the 9,716 topologies capable of producing a sequential advantage. Every sequential topology contains at least one effective positive feedback loop (**Supplementary Figure S4B**), with the distribution peaking at 4–5 loops per topology (**Figure 3C**).

### Bistability underlies the sequential therapy advantage

The universal presence of positive feedback loops across all sequential topologies, established in the preceding analysis, points toward a specific dynamical mechanism: bistability. Positive feedback is a well-established structural prerequisite for generating switch-like, bistable behaviour in biological networks [18–22], and we hypothesised that this property is what enables sequential treatment to outperform concurrent administration.

Our reasoning is as follows. Under combined treatment, the governing equations and kinetic parameters of the system are identical regardless of whether the two drugs were administered concurrently or sequentially - the only difference is the state of the system at the moment combined treatment begins. In a network with a single stable steady state, both administration schedules would therefore converge to the same outcome, and no sequential advantage could exist. For the treatment schedule to determine the final outcome, the network should possess at least two stable steady states under combined treatment, such that the path by which the system arrives at combined treatment - concurrent versus sequential - determines which attractor it ultimately occupies.

To test this hypothesis, we assessed the capacity for bistability across all 127 robust sequential topologies. For each topology, we sampled 100,000 initial conditions under the combined-treatment system and recorded the number of distinct stable steady states reached. Every robust sequential topology exhibited two or more stable steady states, confirming that bistability is a necessary feature of these architectures (**Figure 4A**). The vast majority of topologies displayed exactly two stable states, consistent with the minimal bistable switch expected from a single dominant positive feedback loop. A small number exhibited three stable states, reflecting the additional feedback complexity present in higher-edge topologies.

**Figure 4.**
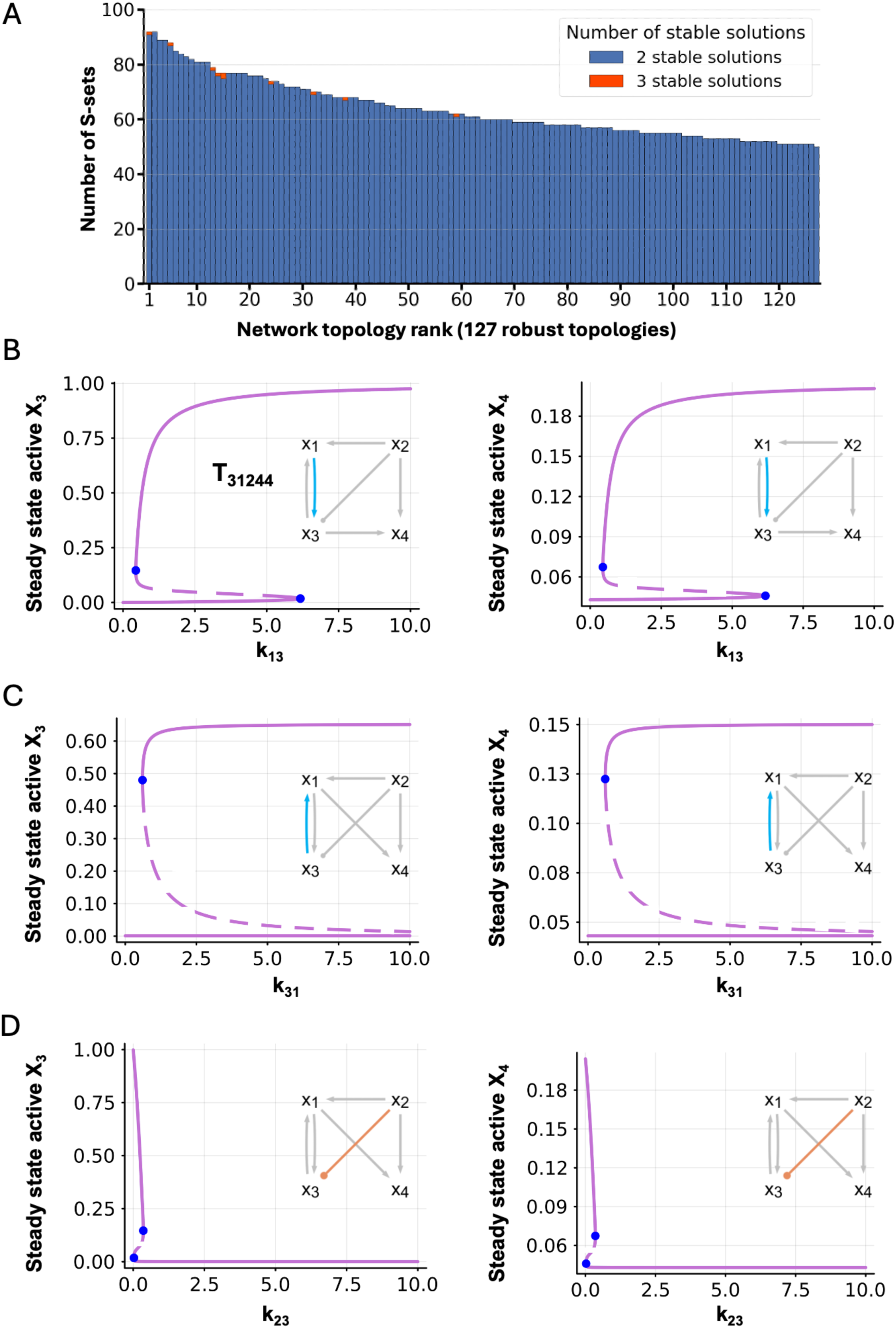
Bistability as the mechanistic basis for sequential therapeutic advantage. **(A)** Distribution of stable steady states across the 127 robust sequential topologies. For each topology, 100,000 initial conditions were sampled under the combined-treatment system and the number of distinct stable states recorded. All robust topologies exhibit at least two stable states (blue), with a small fraction showing three (orange), confirming bistability as a universal feature. **(B–D)** Bifurcation analysis of topology T_31244_ identifying the three critical interactions that generate bistability. Each panel shows the steady-state values of X₃ (left) and X₄ (right) as a function of the varied kinetic parameter, with solid lines indicating stable branches and dashed lines indicating the unstable branch. Insets show the topology with the varied edge highlighted. **(B)** Varying k₁₃ (catalytic rate of edge e₁₃, X₁→X₃): increasing k₁₃ induces a saddle-node bifurcation, transitioning the system from monostability to bistability. **(C)** Varying k₃₁ (catalytic rate of edge e₃₁, X₃→X₁): the reciprocal edge displays an analogous bifurcation, confirming that the mutual activation loop between X₁ and X₃ constitutes the positive feedback required for bistability. **(D)** Varying k₂₃ (catalytic rate of edge e₂₃, X₂→X₃): increasing k₂₃ progressively narrows and ultimately collapses the bistable regime, demonstrating that antagonistic crosstalk from x₂ shapes the boundaries of the bistable landscape.

Having established that bistability is universally present, we next asked which specific network interactions are responsible for generating it. We performed bifurcation analysis using topology T_31244_, one of the two minimal core robust sequential topologies. Because T_31244_ contains fewer feedback structures than the more complex robust topologies, its reduced architecture supports only bistability rather than higher-order multistability, making it an ideal system for isolating the minimal regulatory logic.

By systematically varying individual kinetic parameters, we identified the edges whose perturbation alters the number of stable states in the system. This analysis consistently implicated three interactions as critical determinants of bistability. First, the activating edge from X₁ to X₃ (e₁₃) exhibits a classic bistable bifurcation structure: as its catalytic rate k₁₃ increases, the system transitions from monostability to bistability through a saddle-node bifurcation, generating two distinct stable branches for both X₃ and X₄ (**Figure 4B**). The reciprocal edge from X₃ to X₁ (e₃₁) displays an analogous bifurcation (**Figure 4C**), confirming that the mutual activation loop between X₁ and X₃ provides the positive feedback required for switch-like dynamics, consistent with the established role of such loops in biological memory and cell-fate decisions [18–22]. Third, the inhibitory edge from X₂ to X₃ (e₂₃) proved equally critical: increasing the catalytic rate k₂₃ progressively narrows and ultimately collapses the bistable regime (**Figure 4D**), revealing that antagonistic crosstalk from the second drug target actively shapes the boundaries of the bistable landscape.

Together, these three interactions - a positive feedback loop coupled with antagonistic crosstalk - define the minimal regulatory logic that underpins the network’s capacity to respond differently to sequential versus concurrent perturbations. The mutual activation between X₁ and X₃ generates the bistable switch, while the inhibition from X₂ to X₃ modulates the width of the bistable window, determining the range of kinetic conditions under which two stable states coexist.

This finding provides a mechanistic explanation for the structural requirements identified in the preceding sections. The backbone topology T_31217_, which contains precisely these three interactions, encodes the bistable switch. The additional inter-pathway coupling (e₃₄ or e₁₄) identified through clustering does not generate bistability itself but instead transmits the consequences of the switch to the second oncogenic output, X₄. In this way, the full architecture required for sequential advantage comprises two functionally distinct modules: a bistable switch in the first pathway and a signal relay to the second pathway.

### Gap time defines a critical therapeutic window

Having established that bistability is the mechanistic basis for sequential advantage, we next asked how the treatment schedule determines which of the two stable states the system reaches. Since the combined-treatment equations are identical regardless of administration order, we reasoned the critical variable must be the state of the system at the moment the second drug is introduced. In the sequential regimen, the first drug shifts the system during the gap time, altering the initial condition from which the combined-treatment dynamics evolve. We hypothesise the gap time therefore controls where on the bistable landscape the system begins its trajectory under combined treatment, and thus which attractor it ultimately reaches.

To test this directly, we systematically varied the gap time between D₁ and D₂ administration across robust sequential topologies using topology T_31244_ and measured the resulting steady-state levels of both oncogenic outputs. For X₃, the system remained at a high steady-state level when the gap time was short, indistinguishable from concurrent treatment (**Figure 5A**). Only beyond a critical threshold did the steady-state value drop sharply to a suppressed level, revealing a switch-like transition between the two attractors. The same threshold behaviour was observed for X₄ (**Figure 5B**), confirming that the transition is coordinated across both oncogenic outputs. Above this threshold, the sequential regimen achieves superior suppression compared to concurrent treatment, defining a therapeutic window within which the sequential advantage is realised.

**Figure 5.**
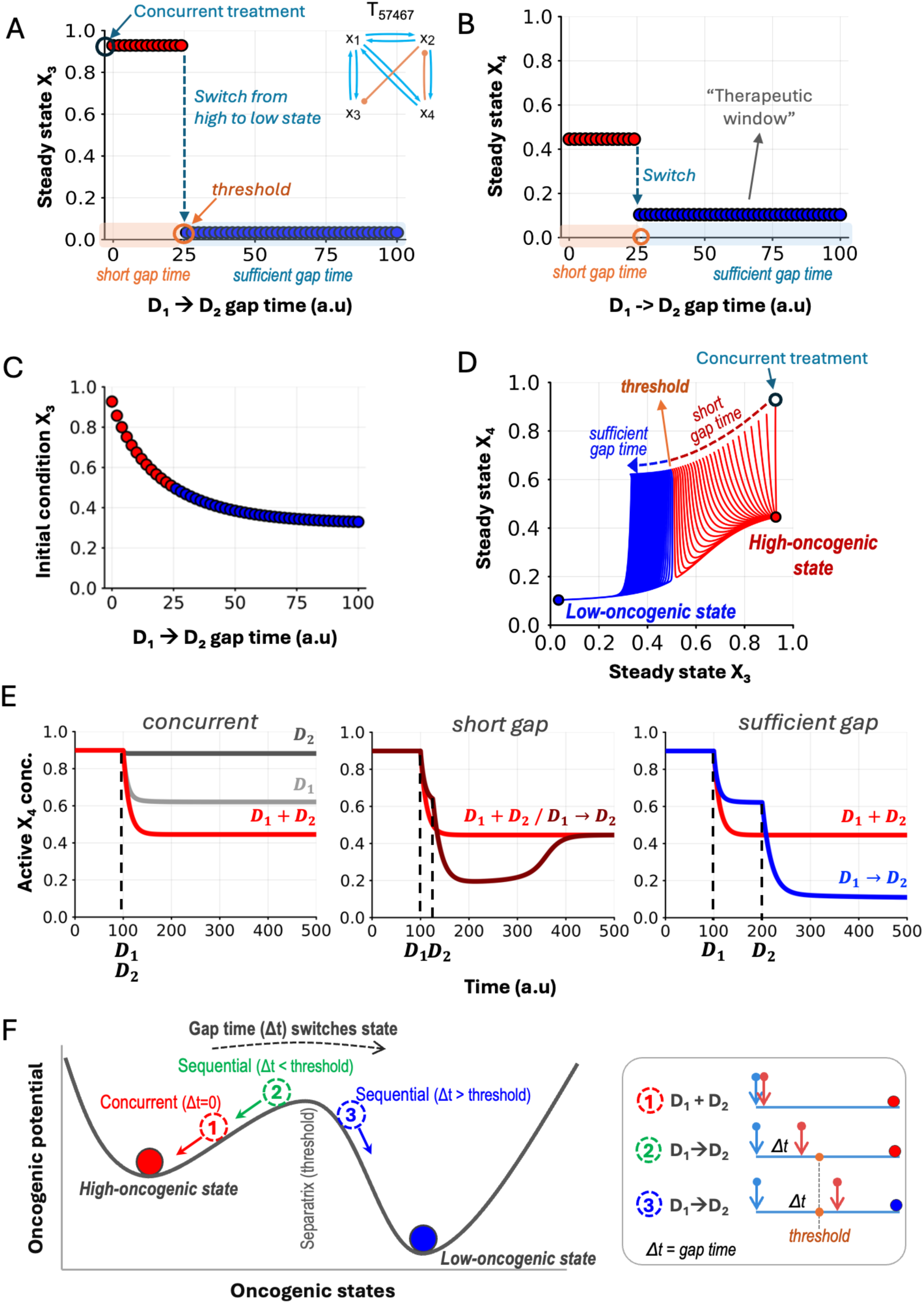
Gap time defines a critical therapeutic window through bistable switching. (A–B) Steady-state levels of X₃ **(A)** and X₄ **(B)** as a function of the D₁→D₂ gap time for topology T_57467_. At short gap times (orange shading), the system remains at a high steady-state level indistinguishable from concurrent treatment (open circle). Beyond a critical threshold (dashed line), both outputs switch sharply to a suppressed state (blue shading), defining a therapeutic window within which sequential treatment achieves superior suppression. **(C)** Initial condition of X₃ at the moment D₂ is introduced, plotted as a function of gap time. As the gap time increases, D₁ progressively suppresses X₃, lowering the initial condition for the combined-treatment system. The colour transition from red to blue marks the crossing of the separatrix between the basins of attraction of the high- and low-oncogenic attractors. **(D)** Phase portrait of X₃ versus X₄ under combined treatment, showing trajectories from multiple initial conditions converging to one of two attractors: a high-oncogenic state (upper right, red) or a low-oncogenic state (lower left, blue). The separatrix divides the basins of attraction. Concurrent treatment (open circle) initiates from the untreated steady state and converges to the high-oncogenic attractor. Sequential treatment with sufficient gap time initiates from a displaced state within the low-oncogenic basin. **(E)** Representative time-course dynamics of X₄ under three treatment scenarios. Left: concurrent administration (D₁ + D₂ at the same time) - the system settles at a high steady-state level. Middle: sequential with short gap time (D₁→D₂, gap below threshold) - the system converges to the same high state. Right: sequential with sufficient gap time (D₁→D₂, gap above threshold) - X₄ is suppressed to the low-oncogenic attractor. **(F)** Conceptual model of the bistable therapeutic landscape. The oncogenic potential landscape under combined treatment contains two stable states separated by a separatrix. Scenario 1 (concurrent, Δt = 0): the system begins in the basin of the high-oncogenic attractor and remains there. Scenario 2 (sequential, Δt < threshold): the gap is insufficient to cross the separatrix, yielding the same outcome. Scenario 3 (sequential, Δt > threshold): the first drug displaces the system past the separatrix, and the combined treatment captures it in the low-oncogenic attractor.

The mechanism underlying this transition is revealed by tracking the state of X₃ at the moment the second drug is introduced, which serves as the effective initial condition for the combined-treatment system (**Figure 5C**). As the gap time increases, the first drug progressively suppresses X₃, pulling the initial condition downward. When the gap time is short, the initial condition remains in the basin of attraction of the high-oncogenic state (red), and the system converges to the same attractor as concurrent treatment. Beyond the critical threshold, the initial condition crosses into the basin of attraction of the low-oncogenic state (blue), and the system is captured by the therapeutically favourable attractor. This crossing corresponds to the steady-state transition observed in **Figures 5A** and **B**.

The full dynamics of this process are visualised in the phase portrait of X₃ versus X₄ under combined treatment (**Figure 5D**). From a range of initial conditions, the system’s trajectories converge to one of two attractors: a high-oncogenic state (upper right) or a low-oncogenic state (lower left), separated by a threshold boundary. Concurrent treatment, which begins from the untreated steady state, invariably converges to the high-oncogenic attractor. Sequential treatment with a sufficient gap time initiates from a displaced state that lies within the basin of attraction of the low-oncogenic attractor, enabling the system to reach a suppressed state that is inaccessible through concurrent administration.

To illustrate how these dynamics unfold in time, we compared the trajectory of x₄ under three representative scenarios (**Figure 5E**). When both drugs are administered concurrently (left panel), the system settles to a high steady-state level. When the second drug is applied shortly after the first (middle panel), the brief gap is insufficient to displace the system past the threshold, and it converges to the same high state as concurrent treatment. Only when the gap time exceeds the critical threshold (right panel) does the first drug shift the system far enough to redirect the trajectory toward the low-oncogenic attractor, achieving sustained suppression that concurrent treatment cannot.

These results are summarised in a conceptual model of the bistable therapeutic landscape (**Figure 5F**). The network under combined treatment possesses two stable states: a high-oncogenic and a low-oncogenic attractor, separated by a separatrix that defines the boundary between their basins of attraction. Concurrent administration (scenario 1) begins from the untreated state, which lies within the basin of the high-oncogenic attractor, and thus converges to the less favourable outcome. Sequential administration with an insufficient gap time (scenario 2) displaces the initial condition but not beyond the separatrix, yielding the same result. Only when the gap time exceeds the critical threshold (scenario 3) does the first drug push the system past the separatrix and into the basin of the low-oncogenic attractor, allowing the subsequent combined treatment to lock the network into a suppressed state.

Overall, these analyses reveal that the gap time is not merely a dosing parameter but a decisive determinant of therapeutic outcome. It defines a critical therapeutic window: too short, and the system behaves as if treated concurrently; sufficiently long, and the sequential regimen accesses a suppressed state that concurrent administration cannot reach. The existence of this window is a direct consequence of the network’s bistable architecture. Without two stable states, there would be no separatrix to cross and no alternative attractor to capture.

## Discussion

### Network design principles underlying sequential advantage

Our systematic enumeration of 59,040 four-node regulatory topologies reveals that the capacity for sequential therapeutic advantage is not a generic property of signalling networks but is strictly constrained by specific architectural features. Of the full ensemble, only 127 topologies robustly produce a sequential benefit, and these share a common structural logic: a positive feedback loop between the primary drug target and its downstream oncogenic output, antagonistic crosstalk from the secondary drug target onto the first pathway, and at least one positive inter-pathway connection that relays the state change to the second oncogenic output.

This minimal architecture can be decomposed into two functionally distinct modules. The first, a bistable switch, is encoded by the mutual activation between X₁ and X₃, whose boundaries are shaped by the inhibitory input from X₂. The second, a signal relay, is provided by a positive edge linking the first pathway to the second oncogenic output X₄, enabling the consequences of the switch to propagate across the full network. The topology atlas analysis further demonstrates that robustness is progressively enhanced through the accumulation of additional positive feedback loops and direct signal propagation routes, establishing a clear hierarchy of structural priorities for sequential efficacy.

These findings recast the sequential advantage, widely observed in empirical studies [11–14] yet lacking a generalisable mechanistic basis, as an emergent, predictable property of network topology. Rather than being an intrinsic property of drug pairs, the potential for sequential benefit is determined by whether the targets of those drugs are embedded within regulatory architectures that support bistability and inter-pathway coupling.

### Bistability as the mechanistic engine of sequential efficacy

The reliance of all robust sequential topologies on positive feedback loops is mechanistically explained by the network’s capacity for bistability. Because the governing equations under combined treatment are identical regardless of the administration schedule, a monostable system would converge to the same steady state irrespective of timing. For a sequential advantage to exist, the network must possess two coexisting stable states, with the treatment schedule serving as the determinant of which attractor is ultimately reached.

Our bifurcation analysis confirms this requirement and identifies the precise interactions responsible. The mutual activation loop between X₁ and X₃ generates the bistable switch through saddle-node bifurcations, while the inhibitory edge from X₂ to X₃ modulates the width of the bistable regime. This interplay between positive feedback and antagonistic crosstalk defines the parametric conditions under which two stable states coexist, and thus the conditions under which sequential treatment can outperform concurrent administration.

In this framework, the network functions as a dynamical memory module: the first drug does not merely attenuate signal intensity but reconfigures the system’s position on the bistable landscape. The subsequent addition of the second drug then locks the network into a suppressed attractor that is inaccessible through concurrent administration alone. This mechanism explains why timing is decisive: the first drug should be given sufficient time to shift the system past the separatrix before the second drug is applied.

While bistability (the coexistence of exactly two stable states) is sufficient to explain the sequential advantage and is the dominant behaviour observed across the robust topologies, a small fraction of more complex architectures exhibit three or more stable states (**Figure 4A**). This higher-order multistability arises from the accumulation of additional positive feedback loops in topologies with greater edge complexity. Although such architectures expand the repertoire of accessible attractors, the fundamental principle remains the same: the treatment schedule determines which attractor the system ultimately occupies. Investigating how multistable landscapes with more than two attractors influence treatment outcomes, including the possibility of intermediate or partially suppressed states, represents an important direction for future work.

### The critical role of gap time in therapeutic outcomes

A direct consequence of the bistable landscape is that the gap time between drug administrations emerges as a critical therapeutic parameter. Our analysis reveals that this interval defines the effective initial condition from which the combined-treatment system evolves: too short a gap, and the first drug has not displaced the system sufficiently far from its original attractor; sufficiently long, and the system crosses the separatrix into the basin of the therapeutically favourable state. The gap time therefore does not just modulate the magnitude of the sequential advantage, it determines whether any advantage exists at all.

This finding carries important implications for clinical dosing. It predicts the existence of a critical therapeutic window, below which sequential and concurrent administration are functionally equivalent, and above which sequential treatment accesses a qualitatively different- and more suppressed - steady state. The sharpness of this transition, observed as a switch-like jump in our simulations (**Figure 5**), suggests that the therapeutic window may be relatively well-defined for a given network architecture and kinetic regime, potentially enabling its prediction from network models.

An important question raised by this analysis is what determines the directionality of the bistable switch, that is, why concurrent treatment converges to the high-oncogenic attractor while sequential treatment reaches the low-oncogenic state, rather than the reverse. In our framework, the pre-treatment steady state lies within the basin of attraction of the high-oncogenic state under combined treatment, and concurrent administration initiates from this state directly. The first drug in the sequential regimen displaces the system toward lower oncogenic activity, and if it crosses the separatrix, the combined-treatment dynamics capture it in the low-oncogenic basin. However, the reverse scenario is equally conceivable for alternative network architectures or parameter regimes: a first drug could, in principle, push the system into a high-oncogenic basin that concurrent treatment avoids. Our selection criterion, requiring that D₁→D₂ produces lower x₃ and x₄ than all other modalities, explicitly filters for the therapeutically favourable direction, but the underlying bistable mechanism is symmetric. Identifying the topological and kinetic features that determine this directionality remains an open question with significant translational relevance.

### Translational Impact and Future Directions

The identification of core topological requirements for sequential efficacy provides a predictive framework for navigating the vast combinatorial landscape of targeted therapies. Our findings suggest that by mapping the regulatory architecture surrounding drug-target pairs, one can assess whether those targets are embedded within network motifs that support bistability and inter-pathway signal propagation - the two prerequisites for sequential advantage. This shifts the paradigm from empirical testing of all possible schedules toward rational selection of candidates whose network context is structurally primed for time-dependent efficacy.

Several concrete applications emerge from this framework. First, the motifs identified here can serve as a predictive catalogue: for a given pair of drug targets, one can assess whether the surrounding regulatory architecture contains the backbone features (positive feedback, antagonistic crosstalk, inter-pathway relay) associated with sequential benefit. Second, the gap time threshold predicted by our analysis suggests that optimal dosing intervals may be estimable from network kinetics, providing a quantitative basis for scheduling decisions. Third, the structural hierarchy revealed by the topology atlas, in which direct signal propagation and X₂ suppression are preferentially retained, may help prioritise among candidate architectures when multiple options exist.

Looking forward, several extensions of this work are possible. Experimentally validating the predicted motifs in well-characterised cancer signalling networks, such as the EGFR–ERK and PI3K–AKT–mTOR axes, where positive feedback loops and pathway crosstalk are well documented, would provide a critical test of the framework’s translational utility. Computational extensions could include scaling the enumeration to larger network sizes, incorporating pharmacokinetic drug dynamics beyond the step-function model used here, and exploring how stochastic fluctuations in signalling activity influence the reliability of bistable switching. Additionally, integrating the topological criteria identified here with patient-specific multi-omics [25,26] data could enable personalised predictions of which tumours are most likely to benefit from sequential dosing.

### Limitations

Several simplifying assumptions in our modelling framework warrant consideration. First, we excluded self-regulatory interactions (autoregulation) to reduce the combinatorial space, yet autoregulation is prevalent in biological signalling networks and could influence both the propensity for bistability and the robustness of sequential advantage. Future analyses that incorporate autoregulatory edges may reveal additional motifs or modify the robustness landscape. Second, the within-pathway activations (X₁→X₃ and X₂→X₄) were fixed as positive throughout the enumeration, which constrains the analysis to architectures in which upstream nodes activate their downstream partners. While this is a common motif in oncogenic signalling, relaxing this constraint could uncover qualitatively different architectural classes.

Third, drug inhibition was modelled as an instantaneous step function with fixed efficacy, abstracting away pharmacokinetic processes such as drug absorption, distribution, and degradation. In reality, drug concentrations rise and decay over time, and the effective gap time experienced by the cell may differ from the administered interval. Incorporating realistic pharmacokinetic profiles would refine the gap time predictions and their clinical applicability. Fourth, our analysis focuses on steady-state outcomes and does not capture transient dynamics that may be therapeutically relevant, such as the duration and depth of signal suppression during the gap period.

Finally, while the four-node framework captures the essential features of two-pathway crosstalk, real signalling networks involve considerably more components, feedback layers, and regulatory complexity. The design principles identified here - positive feedback-mediated bistability, antagonistic crosstalk, and inter-pathway signal relay - represent necessary structural motifs, but their sufficiency in the context of larger, more interconnected networks remains to be established.

## Methods

### Network enumeration

To systematically explore the topological landscape, a four-node network is formulated as a vector of size 16, where each element 𝑒_*ij*_ represents the regulation from node 𝑥_*i*_ to 𝑥_*j*_. There are 3 possible regulation values: 1 (activation), -1 (inhibition), or 0 (null).

An exhaustive analysis of all possible four-node configurations is very computationally expensive and time consuming as it would require the simulation and analysis of 3^16^ = 43,046,721 unique topologies. To mitigate this computational burden and focus on biologically relevant structures, we implemented several constraints. First, we excluded all self-regulatory links (autoregulation), which reduced the system to 12 variable edges. To represent the parallel pathways characteristic of oncogenic signalling, we further fixed edges 𝑒_13_ and 𝑒_24_ to 1. This configuration ensures that upstream proteins consistently activate their respective downstream oncogenic targets.

These constraints effectively focused the enumeration on 10 remaining variable edges, yielding 3^10^ = 59,049 candidate topologies. Finally, we focused specifically on networks that integrate signals between pathways by excluding the 9 topologies where all cross-talk edges (𝑒_12_, 𝑒_14_, 𝑒_21_, 𝑒_23_, 𝑒_32_, 𝑒_34_, 𝑒_41_, 𝑒_43_) were null. This resulted in a final ensemble of 59,040 network topologies for the simulation of various treatment scenarios.

### Mathematical model of network topologies

To simulate the dynamic behaviour of the enumerated topologies under drug treatment, we employed a mathematical framework based on enzymatic networks described by a system of Ordinary Differential Equations (ODEs). Below we clarify the assumptions, definitions, and notations underlying our model construction.

### Key assumptions

I. Network activation occurs through input stimulation at nodes 𝑥_1_ and 𝑥_2_.
II. To balance simplicity and biological realism, we assume the total concentration of each node remains constant and is set to 1. If 𝑥*_i_* represents the active fraction, the inactive fraction is consequently (1 – 𝑥*_i_*). By defining these active fractions as time-dependent variables, we express this conservation as:

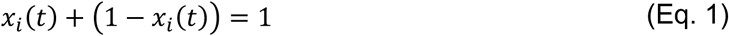

III. A general ODE framework is constructed to describe the behaviour of any 4-node network topology. Several variable and equation parameters are defined for clarity:

- **Topology Parameters:** Vector 𝑒 = (𝑒_11_, 𝑒_12_, …, 𝑒_43_, 𝑒_44_) represents interactions between nodes with each taking values of 1, -1 or 0 as noted previously.
- **Interaction Regulation Parameters:** Vectors 𝑘 = (𝑘_11_, 𝑘_12_, …, 𝑘_43_, 𝑘_44_) and 𝐾 = (𝐾_11_, 𝐾_12_, …, 𝐾_43_, 𝐾_44_) represent the catalytic rates and Michaelis constants, respectively, associate with each regulation 𝑒*_ij_*.
- **Input Stimulus Parameters:** Vectors 𝐼 = (𝐼_1_, 𝐼_2_, 𝐼_3_, 𝐼_4_), 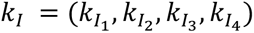 and 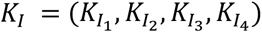 represent external input stimulations at each node, and their corresponding catalytic rates and Michaelis constants respectively.
- **Basal Regulation:** Vector 𝐵 = (𝐵_1_, 𝐵_2_, 𝐵_3_, 𝐵_4_) represents basal enzyme concentration at each node. Associated with these concentrations are binary vectors 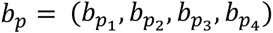 and 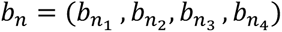 which indicate the presence of the basal activation and degradation respectively. These are couple with their respective rate constants 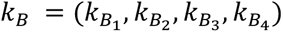 and Michaelis constants 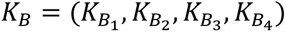. This basal regulation is introduced to prevent the system from unrealistic saturation or depletion of node activity. It ensures each node remains within a responsive biological range by providing continuous opposing regulation. Specifically, if a node lacks positive network-based activation, a basal enzyme provides a constant activating input to prevent depletion. Similarly, if a node is not negatively regulated within the network, a basal enzyme exerts constant inhibition to prevent saturation.
- **Drug Effect Modelling:** Drug inhibition is modelled via the allosteric regulation of the catalytic activities of the drug target node. This is done by effectively lowering the catalytic rate constant 𝑘*_ji_* of all the outbound edges of the node 𝑥*_i_*. We introduce a time-dependent drug function 𝐷(𝑡), expressed as:

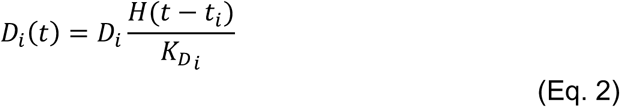

The time-ordered administration of the drug is represented by the Heaviside step function, 𝐻(𝑡 – 𝑡*_i_*), where 𝑡*_i_* is the time of administration. Under this mechanism, the effective drug concentration at node 𝑥*_i_* is defined as 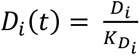 for 𝑡 ≥ 𝑡*_i_* and *D_i_(t)* = 0 for 𝑡 ≤ 𝑡*_i_*. Here, 𝐷*_i_* denotes the administered concentration and 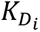 represents the drug inhibition coefficient.

The generalised ODE System for each node is expressed as:

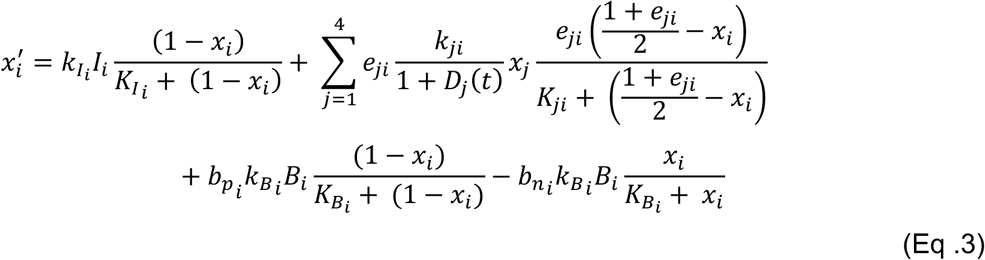

In this equation, the dynamics are governed by four primary components: Term 1 accounts for external input stimulation, while Term 2 defines edge-based regulation within the network. Terms 3 and 4 represent basal activation and basal inhibition, respectively.

### Simulation framework

To comprehensively characterise potential network responses to drug perturbation, we systematically explored the range of dynamic behaviours attainable for each network topology across varying kinetic parameter values. Specifically, we varied the catalytic rates (𝑘*_ij_*) and Michaelis constants (𝐾*_ij_*), which define the strength of interactions between network nodes, while keeping the underlying network topology fixed. 100,000 random parameter sets were generated and used to perform time-dependent in silico simulations to evaluate the dynamic response to drug treatment.

To efficiently sample the parameter space, the Latin Hypercube Sampling (LHS) method was applied. Catalytic rates (𝑘*_ij_*) were sampled within the range 10^-1^ to 10^1^, while Michaelis constants (𝐾*_ij_*) were sampled within the range 10^-2^ to 10^2^. LHS ensures broad and uniform coverage of the parameter space by partitioning each sampling interval into equal segments and selecting one value uniformly from each segment. Random values were initially sampled from a uniform distribution over the interval (0, 1), rescaled to the ranges (−1, 1) or (−2, 2), and subsequently exponentiated to obtain the final parameter distributions corresponding to the desired kinetic ranges.

Following parameter generation, dynamic simulations were performed to evaluate network responses to different drug treatment strategies. For each parameterised topology, an initial pre-treatment simulation was conducted in which the system was stimulated from a small initial value (𝜀) and allowed to evolve in the absence of drug perturbations until a steady state was reached. Steady state convergence was verified by evaluating the gradient of the system and confirming that it approached zero. Stability of the steady state was further assessed by computing the eigenvalues of the Jacobian matrix and confirming that the real parts of all eigenvalues were negative. In addition, state variables were required to remain within biologically meaningful bounds between 0 and 1.

Once the pre-treatment steady state had been established, five treatment scenarios were simulated: monotherapy targeting node 1, monotherapy targeting node 2, concurrent inhibition of nodes 1 and 2, sequential treatment targeting node 1 followed by node 2, and sequential treatment targeting node 2 followed by node 1. Across all network topologies and parameter sets, this resulted in a total of 5 × 59,040 × 100,000 ≈ 30 𝑏𝑖𝑙𝑙𝑖𝑜𝑛 simulations.

Simulation outputs were stored using the Hierarchical Data Format (HDF5) to minimise storage requirements. For each simulation, the topology id, parameter set id and the final steady state concentrations of the nodes: 𝑥₁, 𝑥₂, 𝑥₃, and 𝑥₄ were recorded.

Finally, only topology-parameter set combinations that solved successfully and satisfied the functional criteria across all treatment scenarios were retained for downstream analysis.

All simulations were implemented in Julia using the DifferentialEquations.jl package [27] and executed on the Adelaide University high-performance computing cluster, Phoenix. Parallelisation was achieved using EnsembleThreads, distributing independent simulations across multiple CPU cores in a single-threaded per-core configuration. The stiff system of ordinary differential equations was solved using the Rodas5P solver.

### Topology atlas

The topology atlas was constructed to systematically organise rebound network architectures based on minimal structural differences, allowing the relationships between numerous network topologies to be visualised and analysed. Each topology was encoded as a 16-dimensional signed interaction vector 𝑒, as described previously.

#### Pairwise Hamming Distance

Pairwise structural distances between topologies were quantified using the Hamming distance. The Hamming distance between two topologies counts the number of positions in the interaction vectors that differ, effectively measuring the minimal number of interaction changes required to convert one topology into another.

Formally for two topologies represented by vectors 𝑒 and 𝑒′, the Hamming distance 𝐷(𝑒, 𝑒′) is defined as:

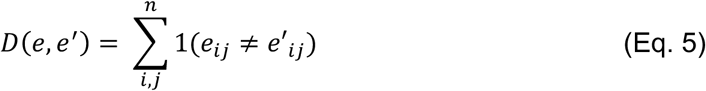

Here, 𝒊 and 𝒋 denote positions within the edge vector, where 𝒊 indicates the source node and 𝒋 indicates the target node.

#### Graph Representation

The atlas is represented as an undirected graph using NetworkX [28] where each node corresponds to a unique topology.

Edges were drawn based on the following criteria:

1. The Hamming distance between two topology vectors equals 1
2. The total number of nonzero interactions differs by exactly 1

This ensures that edges represent minimal structural transformations in which a single interaction is added or removed. Importantly, changing the sign of an existing interaction (activation ↔ inhibition) does not create an edge, since the total number of interactions remains the same. In this way, the graph encodes local neighbourhood relationships between topologies while distinguishing addition/removal of interactions from simple sign flips.

#### Layout

To enhance interpretability, nodes in the atlas were arranged according to structural complexity, represented by the number of interactions in a topology. Nodes were stratified into horizontal layers, with each successive layer representing topologies that have exactly one more interaction than the layer immediately beneath it. For example, nodes in layer 4 contain one additional interaction compared to topologies in layer 3. This vertical stratification creates a clear gradient of structural complexity across the atlas and allows for intuitive visual comparison of networks with different levels of connectivity.

Within each layer, nodes were arranged to minimise edge crossings for visual clarity. Nodes were grouped together according to the lower-layer node(s) they were connected to, so that nodes sharing a connection tend to appear near each other. When a node in the upper layer is connected to multiple nodes beneath it, it is positioned relative to the first node that created the edge. Within each grouping, nodes were further ordered by decreasing robustness, so that the most prevalent topologies appear first. This arrangement reduces visual clutter, highlights clusters of structurally related topologies, and makes paths representing minimal structural transformations easier to follow.

Finally, the coordinate positions of nodes within each layer were equally linearly spaced. In certain layers, however, nodes were manually repositioned to make it easier to trace the paths of connected nodes, highlighting the architectural backbone of topologies.

#### Robustness Visualisation

The robustness of topologies was incorporated directly into the visual attributes of nodes. Node size and opacity were scaled according to a log-normalised frequency metric, such that larger and more opaque nodes correspond to more robust topologies within the dataset. This provides an intuitive way to follow and interpret the variations in robustness across topologies.

To further emphasise specific subsets of topologies, nodes were highlighted using different colours or colourmaps, depending on the criteria or type of topology of interest. The edges connecting these nodes were coloured based on the average of the two connected nodes, providing a visual cue of the relationships between highlighted topologies while preserving robustness information. This approach enables clear visual distinction of important topological features or categories within the atlas.

### Topology Clustering

Network topologies were clustered using hierarchical agglomerative clustering. Pairwise dissimilarity between topologies was computed using the Hamming distance metric (Eq. 5), which quantifies the number of differing edges between binary network representations. Clusters were then formed using the ‘complete’ linkage method, where the distance between clusters is defined as the maximum pairwise distance between their elements.

### Effective Positive Feedback Loop Analysis

To determine whether positive feedback is a universal structural feature of sequential topologies, we quantified the number of effective positive feedback loops present in each of the 9,716 topologies capable of producing a sequential advantage. An effective positive feedback loop is defined as a closed regulatory circuit in which the net sign of the product of all interactions along the loop is positive. This includes both direct positive feedback loops (in which all constituent edges are activating) and double-negative feedback loops (in which an even number of inhibitory edges produce a net-positive circuit).

For each topology, all simple cycles in the corresponding directed graph were enumerated. For each cycle, the sign of the loop was computed as the product of the edge signs along the cycle: a product of +1 indicates a positive (reinforcing) feedback loop, while a product of −1 indicates a negative (balancing) feedback loop. The number of effective positive feedback loops was defined as the count of cycles with a net-positive sign. Cycle enumeration and loop detection were performed in MATLAB using CellNetAnalyzer’s [29] built-in framework, by converting the adjacency list representing the network structure to an incidence matrix.

### Bistability Analysis

To assess the capacity for bistability across all 127 robust sequential topologies, we performed a systematic initial condition screen under the combined-treatment system (D₁ + D₂ applied simultaneously). For each robust topology and its associated parameter sets that exhibited a sequential advantage, we sampled 100,000 initial conditions uniformly across the four-dimensional state space [0, 1]^4^ using Latin Hypercube Sampling. Each initial condition was used as the starting point for a time-integration of the combined-treatment ODE system, which was run until convergence to a stable steady state, verified using the same gradient-norm and Jacobian-eigenvalue criteria described above.

### Bifurcation Analysis

Bifurcation analysis was performed on topology T_31244_ to identify the network interactions responsible for generating bistability. For each edge of interest (*e_13_*, *e_31_*, and *e_23_*), the corresponding catalytic rate parameter (*k_13_*, *k_31_*, or *k_23_*) was varied continuously across the range [0.1, 10] while all other parameters were held fixed at representative values drawn from a parameter set that produces a sequential advantage.

At each value of the bifurcation parameter, the steady states of the combined-treatment system were computed by solving the algebraic equations *dx/dt* = 0 using numerical continuation methods. Stable steady states were identified as solutions for which all eigenvalues of the Jacobian matrix have negative real parts; unstable steady states have at least one eigenvalue with a positive real part. Saddle-node bifurcation points, at which a stable and an unstable steady state coalesce and disappear, were identified as the parameter values where an eigenvalue crosses zero.

Bifurcation diagrams were generated by plotting the steady-state values of *x_3_* and *x_4_* as functions of the bifurcation parameter, with stable and unstable branches displayed as solid and dashed lines, respectively. Bifurcation analysis was implemented in Julia using the BifurcationKit.jl package [30] and the pseudo-arclength continuation (PALC) algorithm, and validated by direct numerical integration from multiple initial conditions at selected parameter values.

### Gap Time Analysis

To investigate how the interval between drug administrations influences therapeutic outcome, we systematically varied the gap time (Δt) between D₁ and D₂ in the sequential regimen D₁→D₂. The analysis was performed on topology T_57467_ using a representative parameter set (S_013322_) that produces a sequential advantage in the primary screen.

#### Gap Time Variation

The gap time was varied from 0 (equivalent to concurrent administration) to 100 arbitrary time units in increments of 1 unit. For each gap time value, the simulation proceeded as follows: (1) the system was initialised at the pre-treatment steady state; (2) D₁ was applied at *t* = 0 and the system was integrated for the specified gap time Δt; (3) at *t* = Δt, D₂ was additionally applied and the system was integrated until convergence to a combined-treatment steady state. The final steady-state concentrations of *X_3_* and *X_4_* were recorded.

#### Threshold Identification

The critical gap time threshold was identified as the value of Δt at which a discontinuous jump in the steady-state levels of *X_3_* and *X_4_* occurs, corresponding to the system crossing the separatrix between the basins of attraction of the high-oncogenic and low-oncogenic attractors. The threshold was determined numerically as the gap time at which the steady-state output first drops below the midpoint between the two attractor values.

#### Phase Portrait and Trajectory Analysis

To visualise the bistable landscape under combined treatment, phase portraits were constructed by simulating the combine treatment system T_57467_- S_013322_ and using a time-sliced snapshot approach. The single-drug (D₁ only) system was simulated from the pre-treatment steady state, and discrete time points were selected from the resulting trajectory. State vectors [*X_1_*, *X_2,_ X_3_*, *X_4_*] at these time points were extracted as snapshots and used as initial conditions for the combined treatment system.

Each initialised state was simulated under dual-drug administration, and trajectories were classified by the attractor they converged to (red, high-oncogenic state; blue, low-oncogenic state). Representative time course plots were generated for three scenarios (concurrent, short gap and sufficient gap) by extracting the trajectory of *X_4_* over a selected range of the full simulation time.

## Data availability

All data generated and analysed in this study are derived from computational simulations and are fully reproducible using the code provided. No experimental datasets were used in this study.

## Code availability

All simulation, analysis, and visualisation code used in this study is available at https://github.com/IntegratedNetworkModellingLab upon request. The repository includes Julia scripts for network enumeration, ODE simulation and bistability/bifurcation analysis; Python scripts for topology clustering, atlas construction, and effective feedback loop analysis; and MATLAB scripts for feedback loop detection.

## Acknowledgements

This work was supported in part by the Australian Research Council (ARC) Centre of Excellence for the Mathematical Analysis of Cellular Systems (MACSYS; CE230100001). This work was also supported in part by a project grant from the National Breast Cancer Foundation, Australia (2023/IIRS0104). The authors thank Dr Ryan Murphy (Adelaide University) and Prof Roger Daly (Monash University) for helpful discussions and critical feedback on the manuscript.

## Author Contributions

L.K.N. conceived and designed the study, developed the conceptual framework, led the data interpretation and analysis strategy, supervised all aspects of the project, and secured funding. T.O.O. and K.I.R. performed all computational modelling, simulation, data processing, and analysis under the supervision of S.-Y.S. and L.K.N. A.H. co-supervised T.O.O., provided critical feedback on the analytical approach and manuscript drafts, and contributed to the interpretation of results. T.O.O. and L.K.N. wrote the manuscript with input from all authors.

## Supplementary Figures

**Supplementary Figure S1.**
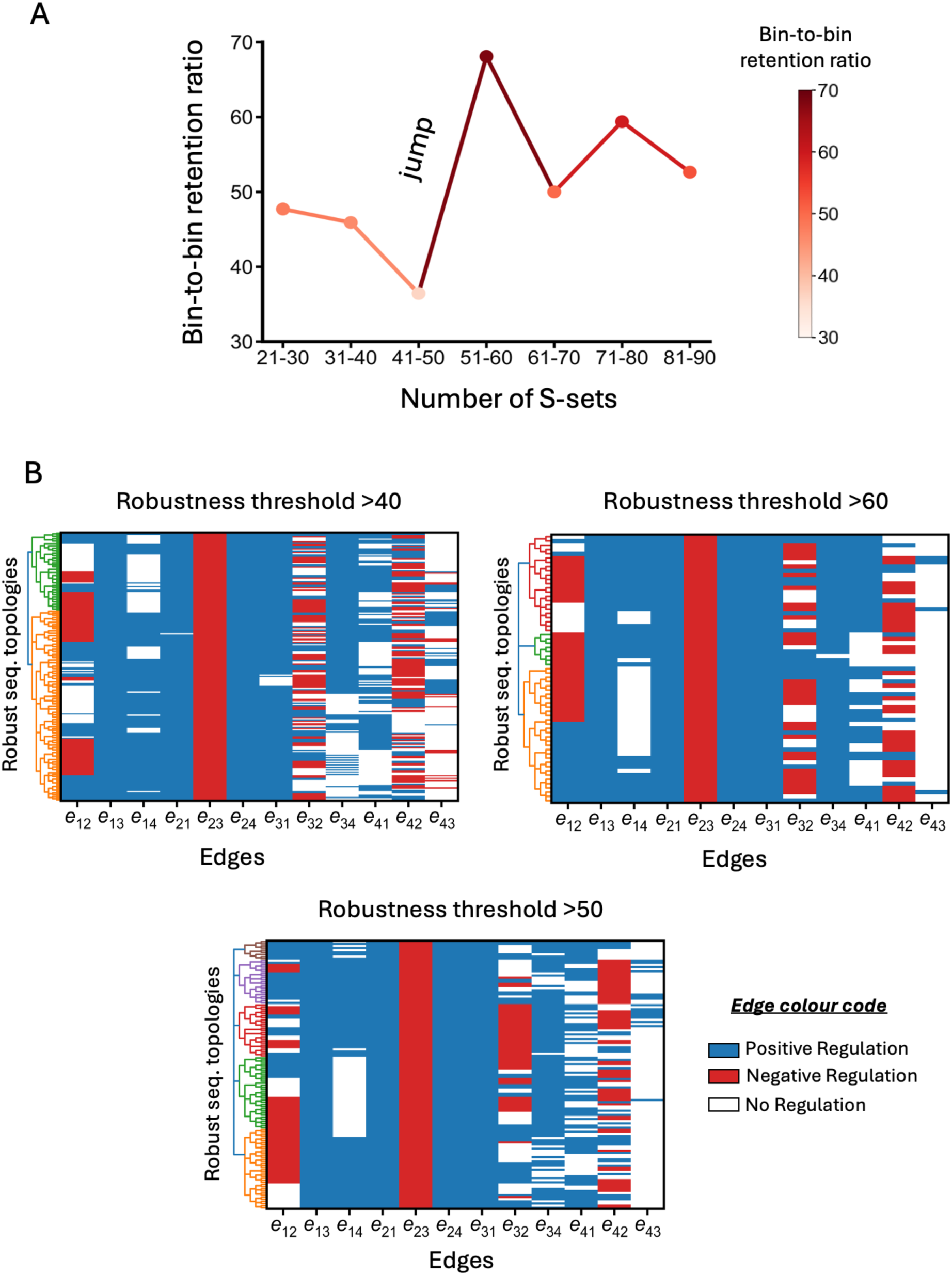
Robustness threshold selection and sensitivity analysis. **(A)** Bin-to-bin retention ratio across sequential set frequency bins. For each bin (in intervals of 10 sequential sets), the retention ratio quantifies the proportion of topologies that persist into the next higher bin. A sharp inflection at approximately 50 sequential sets indicates that topologies above this threshold represent a structurally distinct subset, providing the rationale for the robustness cutoff used in the main analysis. **(B)** Sensitivity of structural findings to the robustness threshold. Clustering and interaction prevalence analyses repeated at thresholds of 40, 50, and 60 sequential sets yield largely consistent backbone architectures and cluster structures, confirming that the identified design principles are stable properties of the sequential topology landscape.

**Supplementary Figure S2.**
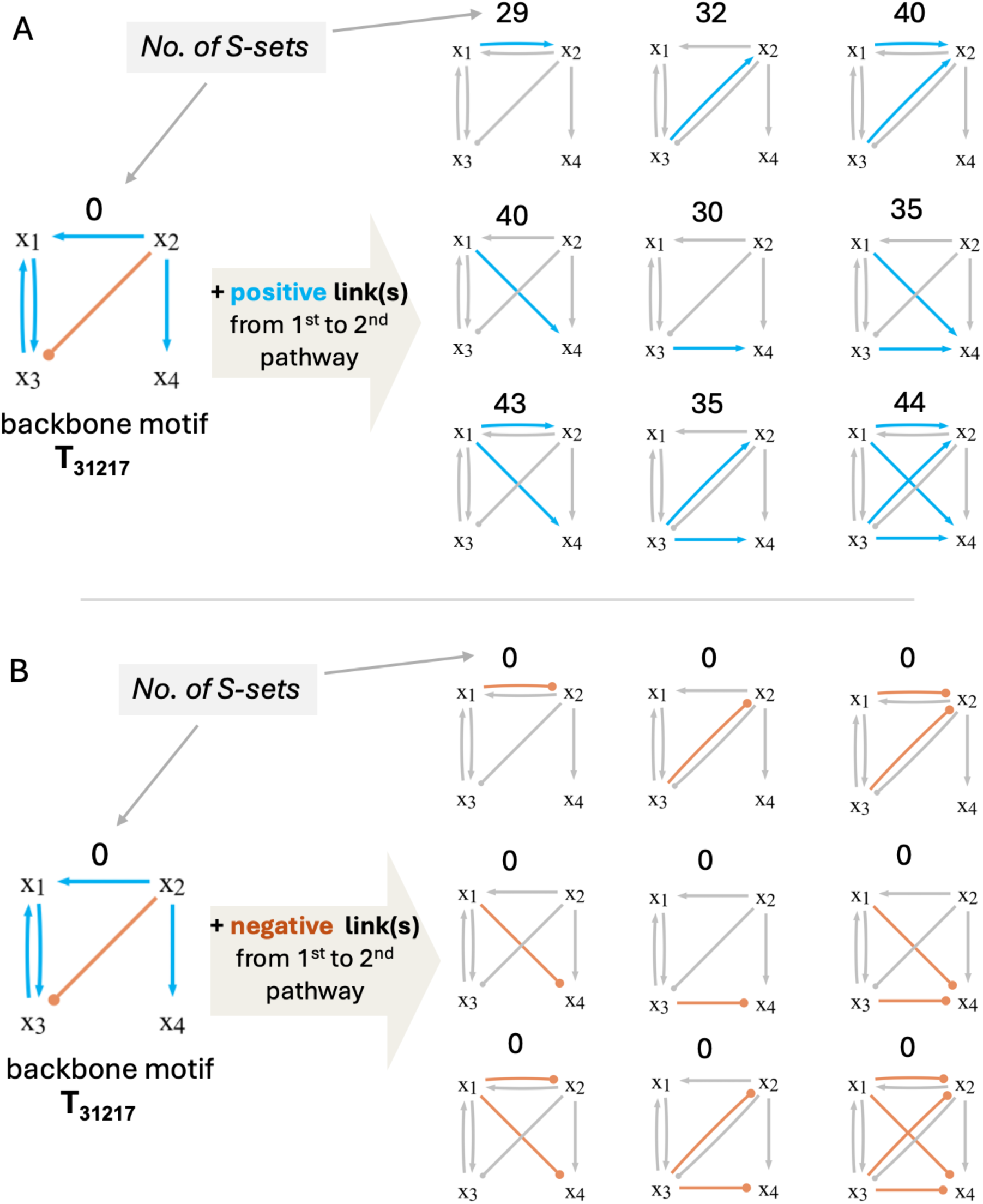
Validation of the inter-pathway coupling requirement. Systematic enumeration of backbone topology T_31217_ variants with all possible combinations of positive, negative, and null connections from X₁ and X₃ to X₂ and X₄. Top: variants with a positive inter-pathway connection produce sequential advantage (non-zero sequential set counts indicated). Bottom: variants with negative inter-pathway connections uniformly fail to produce sequential behaviour, confirming that positive signal propagation from the first pathway to the second is a necessary structural requirement.

**Supplementary Figure S3.**
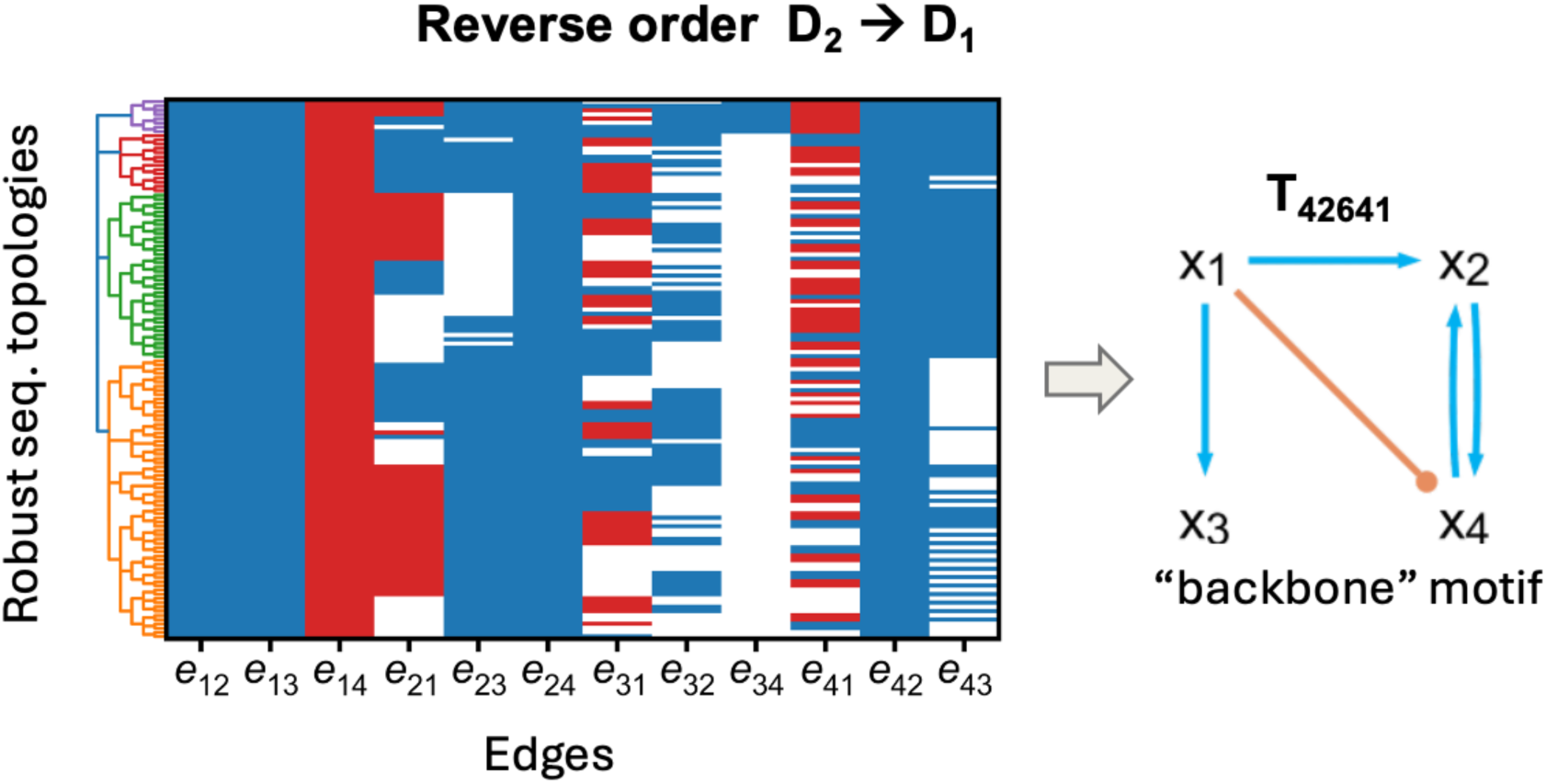
Structural symmetry of the reverse sequential order (D₂→D₁). Clustering and interaction prevalence analysis of robust topologies identified under the D₂→D₁ sequential criterion identifies the structurally symmetric backbone T_42641_, confirming that the architectural requirements for sequential advantage are mirrored across drug orders.

**Supplementary Figure S4.**
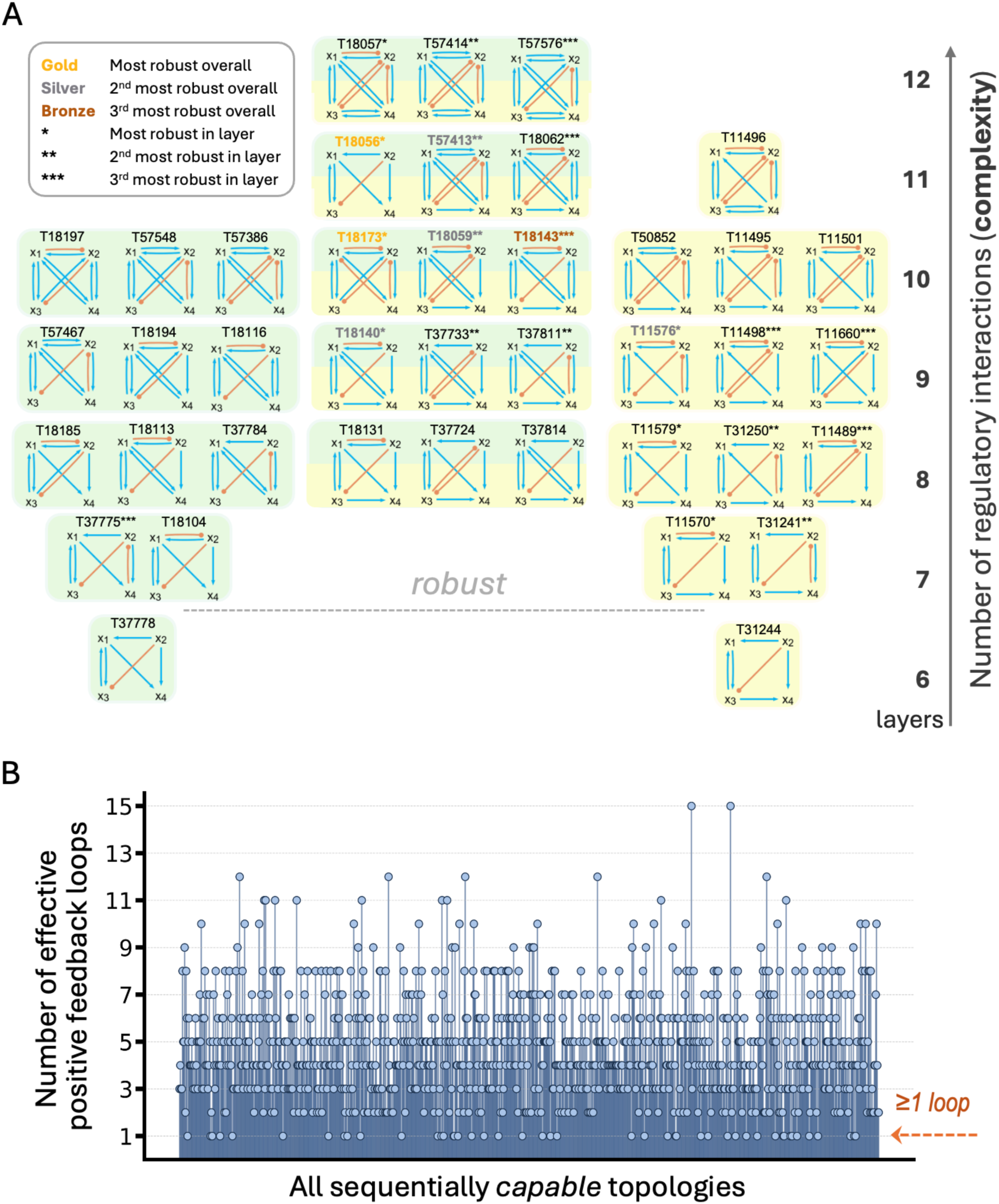
Complete structural progression and effective positive feedback loop analysis. **(A)** Structural progression of the two dominant families across increasing complexity. Topologies derived from T_37778_ (characterised by edge 𝑒_14_) are shown in green, those derived from T_31244_ (characterised by edge 𝑒_34_) are shown in yellow, and topologies containing features of both families are shown in green–yellow. Representative topologies are shown for each layer, illustrating how additional interactions give rise to robust sequential behaviour. Gold (T_18056_, T_18173_), silver (T_57413_, T_18059_), and bronze (T_18143_) denote the overall most, second most, and third most robust topologies, respectively. Asterisks indicate the relative robustness ranking within each layer, with one asterisk representing the highest-ranking topology, followed by increasing numbers for lower ranks. **(B)** Number of effective positive feedback loops (including both direct positive feedback and double-negative feedback) per sequential topology. Each bar represents one of the 9,716 topologies capable of producing a sequential advantage (x-axis), with the y-axis indicating the number of effective positive feedback loops identified. The distribution complements the histogram in Figure 3C by showing the per-topology variation. Every sequential topology contains at least one effective positive feedback loop.

